# Light adaptation explains different linearity of ON and OFF responses

**DOI:** 10.1101/039248

**Authors:** Matteo Carandini

## Abstract

The visual system separates the responses to dark and light stimuli into OFF and ON channels. Recordings in visual thalamus indicate that these channels exhibit marked asymmetries in linearity: OFF responses grow roughly linearly with intensity decrements, but ON responses can saturate with small intensity increments. Here I show that this asymmetry is likely to originate in photoreceptors, as it follows from the classical description of their light responses and of their properties of light adaptation. Under this interpretation, the surprising aspect of the recordings is that the visual system does so little to change this asymmetry in subsequent stages.

## Results

The visual system takes the first opportunity, right after the photoreceptors, to separate the responses to stimuli darker and lighter than the background onto OFF and ON channels. These channels progress separately from retina on to lateral geniculate nucleus (LGN) and thence to cortex. Studies of the visual system have explored the differences between these channels. One of these studies [1], specifically, demonstrated marked differences in linearity, reporting that “OFF neurons […] increase their responses roughly linearly with intensity decrements, independent of the background intensity”, whereas “ON neurons saturate their responses with small increases in intensity and need bright backgrounds to approach the linearity of OFF neurons. “. Various observations in this study point to the phenomenon originating in the photoreceptors, i.e. in pronounced asymmetries in photoreceptor responses to light increments and decrements. Are such asymmetries likely?

Consider the classical description of photoreceptor response *R* to light intensity *I* established by Naka and Rushton in 1966 [2] (Figure 1 **A**, reviewed in Ref. [3]):

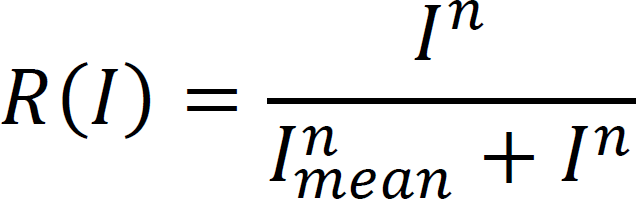

where the exponent *n* is a fixed parameter (here set to 1.37), and *I_mean_* is the mean light intensity. This expression captures light adaptation because the response curve shifts horizontally on a logarithmic axis to remain centered on mean intensity (Figure 1 **A**). When light spots are flashed on a black, gray, or white background (intensities of 1, 50, or 100), the curves center on mean intensities of, say, 5, 50, and 95 (mean intensities are slightly more gray than backgrounds, as only light spots are used on the dark background, and vice versa).

**Figure 1.**
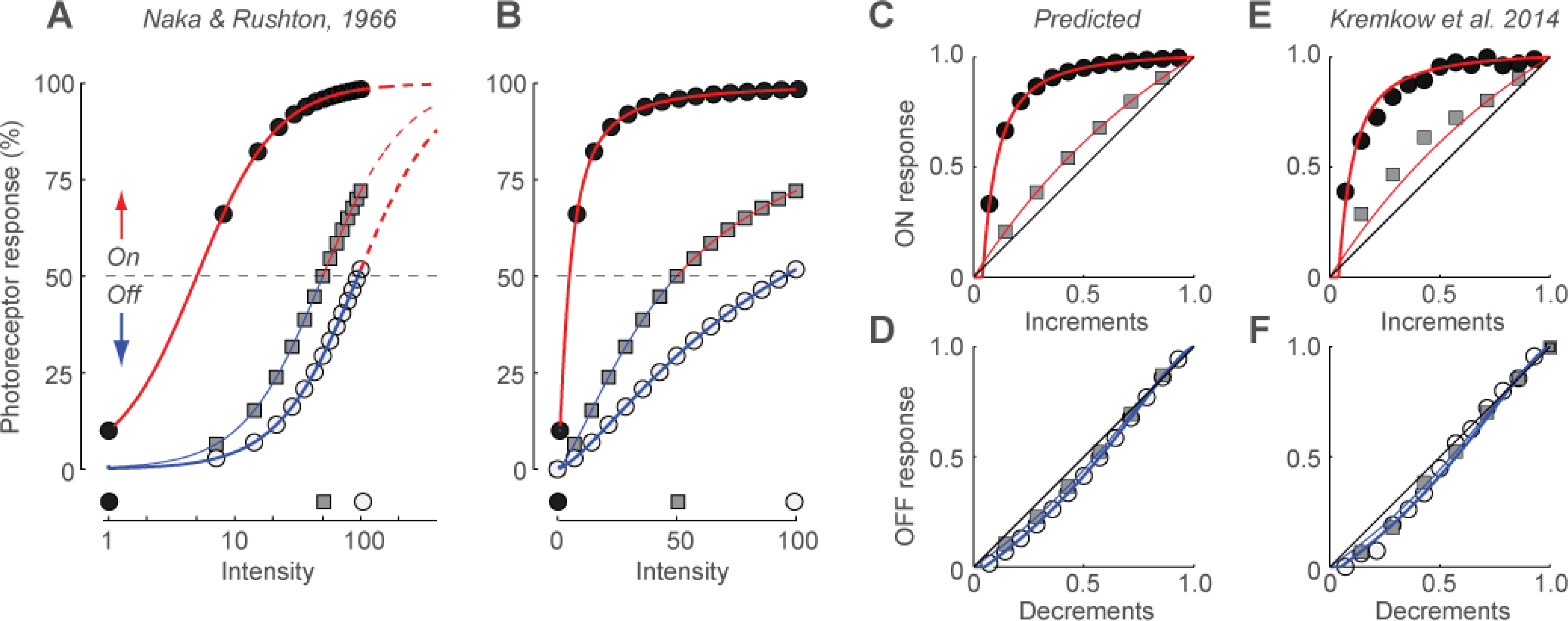
Light adaptation explains differences between ON and OFF responses. **A**. Photoreceptor responses expected when the background is black, gray, or white, in response to light increments (red) or decrements (blue). **B**. The same responses, plotted on a linear axis, up to the maximum screen intensity (notionally, 100). **C**: The responses to light spots (ON responses) presented on a black background (thick) or on a gray background (thin). **D**: The responses to dark spots (OFF responses) presented on a white background (thick) or on a gray background (thin). **E,F**: The same curves, plotted on top of the data measured in LGN cells, replotted from Fig. 3A-D in Ref. [1]. Data were obtained graphically with the Matlab function “grabit”.

When plotting these responses in linear scale, one readily notices a marked asymmetry in the responses to light decrements and increments (Figure 1 **B**). Responses to light increments, shown in red, saturate more strongly than responses to light decrements, shown in blue. These differences would be readily visible in the responses of OFF cells and ON cells (Figure 1 **C,D**). OFF responses grow roughly linearly with intensity decrements, whether the background is gray or white (Figure 1 **D**). ON responses, instead, saturate with small increases in intensity when the background is black, and need a gray background to approach the linearity of OFF responses (Figure 1 **C**). This is very close to the behavior observed in ON and OFF LGN cells (Figure 1 **E,F**, replotted from Ref. [1]). Those observations, therefore, seem largely predicted by light adaptation at the photoreceptors.

This result provides a firm basis to the interpretation given in the original study [1]: the marked differences in linearity between OFF and ON responses are likely to be due to pronounced asymmetries in photoreceptor responses to light increments and decrements. Indeed, the analysis above shows that such asymmetries are fully to be expected based on textbook notions of light adaptation. The surprising aspect of the experimental data [1], therefore, is perhaps not that there is such an asymmetry, but rather that the early visual system past the retina does so little to change it, leaving it as pronounced as it is in the photoreceptors.

## Acknowledgments

MC holds the GlaxoSmithKline / Fight for Sight Chair in Visual Neuroscience.

## References

1 Kremkow, J., et al, Neuronal nonlinearity explains greater visual spatial resolution for darks than lights. Proc Natl Acad Sci U S A, 2014. 111 (8): p. 3170–5.

2 Naka, K.I. and W.A. Rushton, S-potentials from luminosity units in the retina of fish (Cyprinidae). J Physiol, 1966. 185 (3): p. 587–99.

3 Carandini, M. and D.J. Heeger, Normalization as a canonical neural computation. Nat Rev Neurosci, 2012. 13 (1): p. 51–62.

